# Simultaneous Label-free Imaging of Nucleolar Dynamics and Subcellular Metabolic Shifts Across Tissue Contexts

**DOI:** 10.1101/2025.10.07.681013

**Authors:** Kunzan Liu, Ellen L. Kan, Honghao Cao, Jiashu Han, Giovanni de Nola, Linda G. Griffith, Eliezer Calo, Sixian You

## Abstract

The nucleolus is essential for ribosome biogenesis and for regulating cellular responses to growth and stress, and its relationship to cellular metabolic activity and functional state highlights its potential as a biomarker of cellular health. However, challenges in contrast multiplexing and high-resolution isotropic three-dimensional (3D) imaging hinder the non-invasive, simultaneous assessment of nucleolar activity and subcellular metabolic maps across different tissue contexts, especially in complex 3D environments. To fully harness the nucleolus’s potential as a biomarker and diagnostic target, we present a multimodal imaging platform that combines third harmonic generation (THG) imaging with metabolic autofluorescence of NAD(P)H and FAD to study structural and metabolic nucleolar dynamics. Enabled by a high-power multimode fiber source and an axial deblurring network, we achieved ∼ 400 nm isotropic resolution in deep 3D imaging and confirmed the high accuracy of our method for label-free nucleolus identification using co-registered immunostaining and electron microscopy. To establish the biological relevance of our approach, we demonstrate that nucleolar stress leads to an unexpected depletion of NADH across cellular compartments. Furthermore, in the human endometrium—where nucleolar dynamics are central to the tissue’s response to progesterone—our label-free imaging strategy delineated endometrial structures in freshly excised tissues and revealed that progesterone treatment induces distinct changes in nucleolar translocation and metabolic adaptation in organoids derived from diseased patients compared to controls. This capacity to non-invasively visualize and quantify features of the nucleolus and its local metabolic microenvironment at single-cell resolution in human tissues—and dynamically track these changes over time in patient-derived organoids—provides a powerful tool for uncovering the roles of the nucleolus in development, disease progression, and therapeutic response. Together, these findings establish our platform as a significant advance for both fundamental research and organelle-based tissue diagnostics.

## Introduction

The nucleolus is a prominent, membraneless organelle within the cell nucleus that plays a central role in ribosome biogenesis (1). Ribosomes are essential for protein synthesis, and because ribosome production is closely linked to cellular growth and proliferation, nucleolar activity dynamically adjusts in response to metabolic demands—making it a sensitive indicator of the cell’s physiological state. For instance, cancer cells often display enlarged nucleoli and elevated ribosome synthesis, reflecting high proliferative activity (2), whereas quiescent stem cells, with low nucleolar activity, show reduced translational output (3). Changes in nucleolar activity are frequently accompanied by alterations in its structure, shape, number, and localization within the nucleus (4–6). This is especially notable when the nucleolus is functionally inactivated, such as during genotoxic stress responses (7–9), or in hormonally regulated tissues like the endometrium, where nucleolar remodeling and repositioning can reflect cyclical tissue regeneration (10–12). Such spatiotemporal alterations in nucleolar structure and function can provide valuable insights into both normal physiological processes and pathological states, making the nucleolus a promising target for cellular state monitoring across diverse tissue types (13–20).

Although changes in nucleolar structure have been correlated with cell metabolism, direct measurements linking nucleolar state to overall cell state have not yet been formally established. This knowledge gap is in part due to the fact that current imaging technologies fall short in enabling non-invasive assessment of nucleolar activity, limiting our ability to determine how nucleolar function relates to cell state. Most existing approaches rely on label-based methods using fluorescently tagged antibodies, which offer high specificity for detecting nucleolar components but require cell fixation (21). This restricts observations to non-living samples and prevents the study of dynamic processes. While fluorescent protein tagging enables live-cell imaging and addresses this limitation (22), it can interfere with native protein function and is often difficult to apply in primary cells or complex tissue samples such as biopsies.

A fundamental challenge in imaging the nucleolus lies in its biological complexity. As a dense organelle composed of hundreds of proteins and various RNA species (23, 24), labeling only one or a few molecular components offers an incomplete view of its dynamic organization and function. To overcome this limitation, label-free imaging approaches are increasingly being explored. These techniques leverage intrinsic material properties, such as electron density or refractive index, to visualize the nucleolus without the need for external labels, enabling more holistic and exploratory investigations. For example, electron microscopy provides ultra-high-resolution images of nucleolar architecture for detailed structural analysis (25). However, its requirement for ultrathin, fixed samples and the inherently low throughput limit its utility in studying dynamic processes or cellular heterogeneity across large populations. Alternatively, phase-contrast and quantitative phase imaging enhance contrast in transparent samples and are widely used to observe nucleolar dynamics in live cells under accessible setups (26–28). Still, they are limited to transparent 2D samples, making them unsuitable for intact tissues or complex three-dimensional (3D) cultures. A recent advancement introduced laser-emitting cytometry for nucleolar imaging, offering improved contrast and sensitivity over conventional phase-contrast methods, though sample thickness remains a limiting factor (29). Third harmonic generation (THG) microscopy has also been shown as a viable method for detecting the high refractive index of the nucleolus by capturing signals at the water-protein interface (30–32). With the use of near-infrared excitation and intrinsic optical sectioning, THG has the potential for deeper tissue imaging (33, 34), but its adoption in biological research is limited by the low specificity of its contrast mechanism, minimal validation in complex systems, and challenges in interpreting signals arising from multiple coexisting sources.

Here, we present a multimodal imaging platform that combines depth penetration, high 3D spatial resolution, and metabolic sensitivity to fully harness the nucleolus’s potential as a biomarker and diagnostic target. Leveraging the higher refractive index of the nucleolus (*n* = 1.375–1.385 (35)) relative to the rest of the nucleus (*n* = 1.355–1.365 (35)), we structurally identify the nucleolus with THG signals and simultaneously map subcellular metabolism from endogenous NAD(P)H and FAD fluorescence (36–40). This simultaneous multimodal imaging is achieved by simultaneously exciting multiharmonic signals and autofluorescence (34, 41), enabled by a custom-engineered high-power excitation beam in the fundamental mode from a multimode fiber through fundamental mode launching, spectral filtering, and fiber shaping, which enhances both deep light penetration and high spatial resolution. To further improve the axial resolution, we developed SSAI-3D-v2 (System- and Sample-Agnostic Isotropic 3D Microscopy (42), version 2), a deep learning-based axial deblurring network trained on ∼10 GB of diverse structural and metabolic imaging data. SSAI-3D-v2 demonstrates state-of-the-art performance by minimizing domain shift, with expanded applicability across cell types and conditions. With coregistered validation using immunostaining and electron microscopy across multiple cellular systems, we confirmed 96% accuracy in label-free nucleolus identification and achieved ∼400 nm isotropic resolution in 3D. We validated the platform in live cells under both physiological and stress conditions, as well as in intact human endometrial tissue and 3D patient-derived endometrial organoids. As a proof-of-concept for the application of this technology in disease contexts, we present a case study demonstrating that progesterone treatment gives rise to prominent differences in nucleolar translocation and metabolic adaptation in endometrial organoids derived from diseased patients versus control donors. This ability to perform noninvasive visualization and quantification of nucleolar dynamics with subcellular resolution—while simultaneously determining local metabolic state—offers a powerful tool to investigate the role of the nucleolus in development, disease progression, and therapeutic response, establishing this platform as a significant step forward in the advancement of organelle-based tissue diagnostics.

## Results

### Three-dimensional high-resolution structural and metabolic imaging of the nucleolus

To achieve high-resolution 3D structural and metabolic imaging of the nucleolus, we developed a label-free high-resolution multimodal system aided by an improved excitation source and an axial deblurring algorithm (Fig. 1a-c). First, multimodal structural and metabolic imaging was achieved by integrating structural information (THG for optical interfaces and second harmonic generation (SHG) for fibrillar collagen (43, 44)) and metabolic information (autofluorescence from metabolic coenzymes NAD(P)H and FAD) within a microenvironment through a single excitation at 1100 nm (41) (Fig. 1a; see Supplementary Fig. S1 for optical setup). The fidelity of these metabolic readouts was validated *in vitro* using NAD(P)H and FAD standards, yielding the expected nonlinear power dependence consistent with multiphoton generation efficiency, and in cultured cells treated with daporinad, a potent nicotinamide phosphoribosyltransferase (NAMPT) inhibitor, which caused a clear reduction in NAD^+^ levels (Supplementary Fig. S2; see Methods for experimental details).

**Fig. 1:**
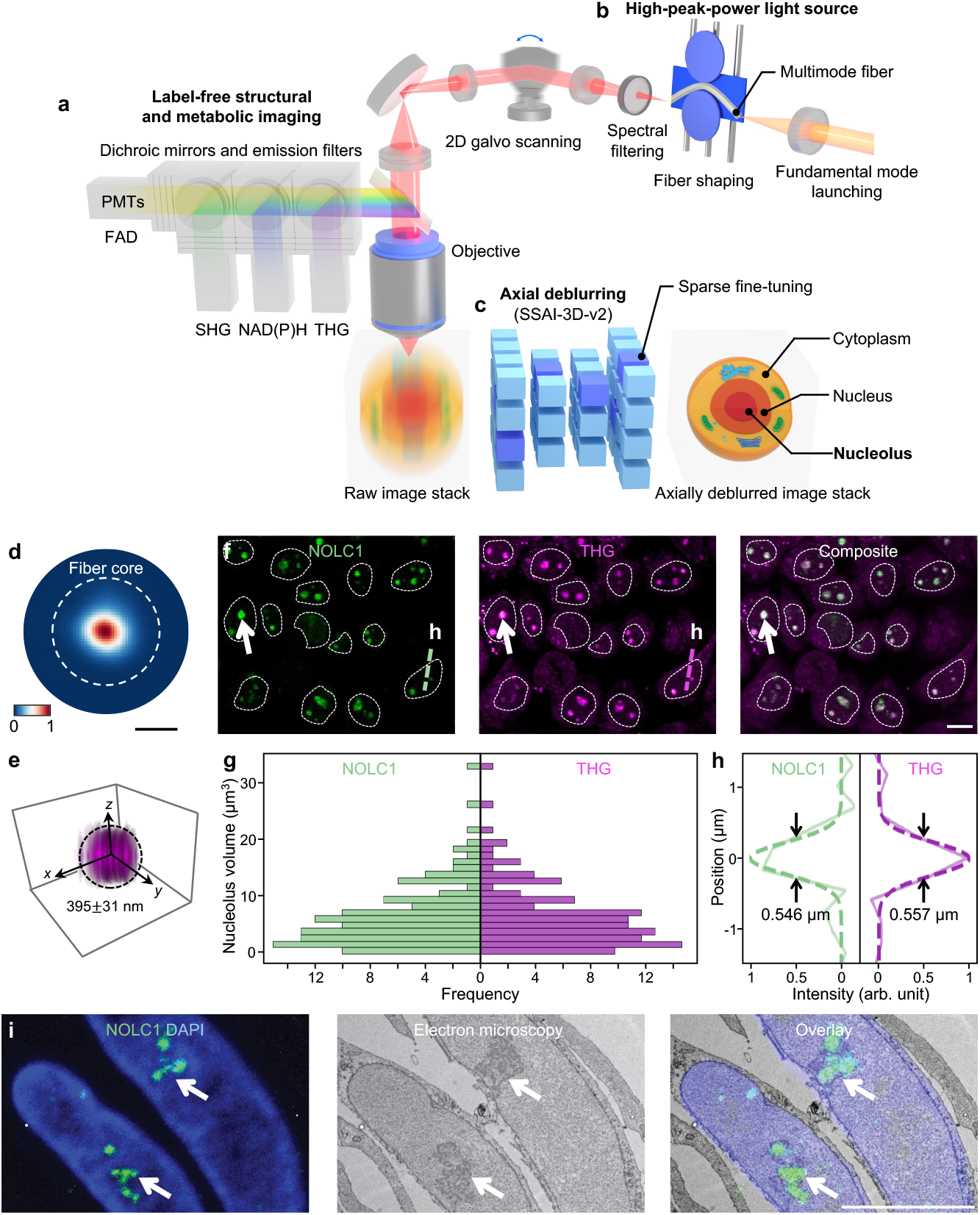
Three-dimensional high-resolution structural and metabolic imaging of the nucleolus. **a**, Label-free structural and metabolic imaging with integrated signals from harmonic generation (SHG and THG) and autofluorescence (NAD(P)H and FAD). **b**, High-power excitation source in the fundamental mode is generated through multimode fiber by integrating fundamental mode launching, spectral filtering, and fiber shaping. **c**, Axial deblurring network for 3D isotropic resolution using SSAI-3D-v2. **d**, The beam profile used for imaging. The white dotted circle represents the core of the multimode fiber. **e**, 3D rendering of THG imaging of sub-diffraction-limited microspheres after axial deblurring. **f**, Validation of the nucleolus by immunofluorescence with the nucleolar marker NOLC1 in HeLa cells. Left: NOLC1. Middle: THG. Right: composite image. White arrows: nucleolus. White dotted regions: nucleus. **g**, Statistics of nucleolus volume derived from NOLC1 and THG. **h**, Line intensity profile of NOLC1 and THG in **f. i**, Validation of the nucleolus in HeLa cells using correlative light and electron microscopy (CLEM). Left: NOLC1 and DAPI. Middle: electron microscopy. Right: overlay image. White arrows: nucleolus. Scale bars: 10 µm.

The use of a near-infrared excitation wavelength allows for deeper imaging by reducing light attenuation (34), while the efficient excitation and acquisition of multimodal signals enable extended time-lapse dynamic imaging with permissible laser power (41). We optimized the excitation source using our previous multimode fiber and fiber shaper design (34, 45), achieving a fundamental mode output beam (Fig. 1d) that provides a lateral imaging resolution of ∼400 nm and an axial resolution of ∼1000 nm (Fig. 1b; mechanisms are discussed in Supplementary Note 1 and Supplementary Fig. S3). Secondly, to enhance axial resolution, we adapted our previous training framework, SSAI-3D (42), and developed and released an improved large-scale axial deblurring network, SSAI-3D-v2. The original SSAI-3D framework aimed to mitigate domain shift in axial deblurring by artificially blurring lateral images to customize a pre-trained network for a given image stack. In contrast, SSAI-3D-v2 further enhances axial deblurring fidelity by leveraging an expanded training dataset and minimizing domain shift through the inclusion of historically diverse 2D images (Fig. 1c; mechanisms are discussed in Supplementary Note 2 and Supplementary Fig. S4 and S5). The improved SSAI-3D-v2 enables isotropic resolution recovery, significantly enhancing axial resolution to match lateral resolution (Fig. 1e).

To confirm that this 3D high-resolution structural and metabolic imaging allows us to resolve and identify the nucleolus, we conducted a validation study in HeLa cells by performing immunofluorescence staining for the nucleolar protein NOLC1 (46, 47) (Fig. 1f). Co-registration of THG signals with fluorescence from NOLC1 staining confirmed the capability of THG for nucleolar imaging, with signals detectable at the water–protein interface (31) (Fig. 1e). The consistency in nucleolus volume statistics and intensity line profiles derived from the two imaging modalities further demonstrates the sensitivity of THG for nucleolar visualization (Fig. 1g and h). We also performed correlative light and electron microscopy (CLEM) in HeLa cells to verify that NOLC1 staining faithfully marks nucleoli (Fig. 1i).

### Disruption of nucleolar function elicits broad, system-wide metabolic flux spanning multiple cellular compartments

To demonstrate our platform’s ability to capture quantitative spatiotemporal changes in metabolic flux during nucleolar perturbation, we first applied it to live cells exposed to stressors known to alter nuclear structure and function (Fig. 2a). Our goals were twofold: to validate that our technology captures known aspects of the nucleolar response to previously characterized perturbations, and to generate new insights by combining structural and metabolic imaging that track spatial and temporal changes following nucleolar stress.

**Fig. 2:**
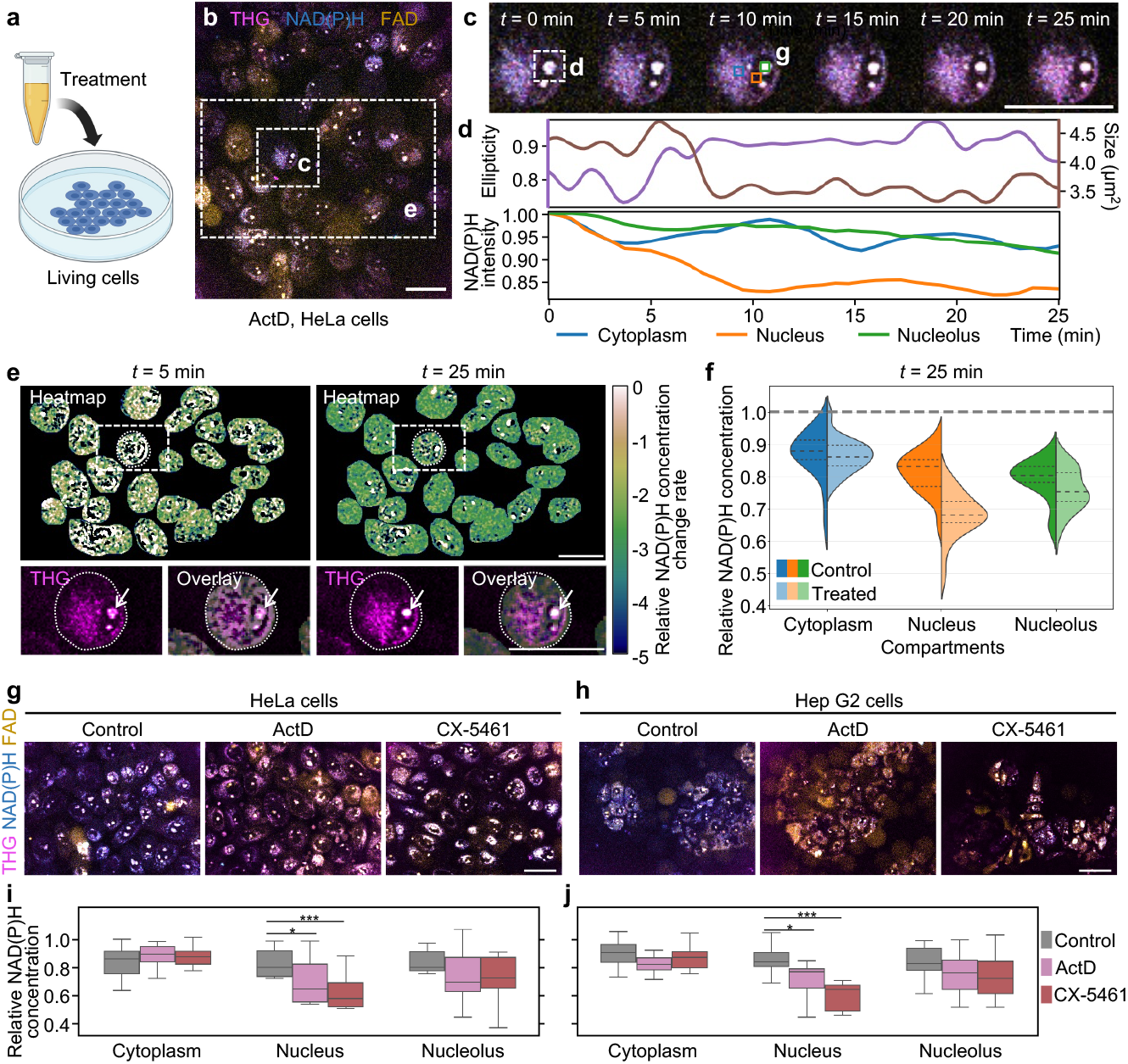
Spatiotemporal nucleolar dynamics in cells in response to perturbations. **a**, Experimental setup of unlabeled living cells with different treatments. **b**, 25-minute time-lapse composite structural and metabolic imaging at 1 Hz. White dotted boxes: regions of the zoomed-in analysis in **c** and **e. c**, Structural and metabolic behaviors of a representative cell over 25 minutes. **d**, Structural (top) and metabolic (bottom) dynamics during the recordings of the nucleolus in **c**, including changes in nucleolar ellipticities and sizes, and changes in NAD(P)H intensity in the cytoplasm, nucleus, and nucleolus. **e**, Pixel-level heatmap of the relative NAD(P)H concentration change rate at *t* = 5 min and *t* = 25 min. White dotted boxes: a cell outlined with a white dotted curve, showing the THG image and the THG image overlaid with the heatmap. White arrows: the same nucleolus in **c** and **d**, highlighting the associations of structural and metabolic changes. **f**, At *t* = 25 min, the relative concentration of NAD(P)H compared to *t* = 0 min in the cytoplasm, nucleus, and nucleolus. Control and treated indicate experiments without and with ActD treatment, respectively (*n* = 20 cells for each violin plot). **g** and **h**, Extended experiments with different treatments (ActD and CX-5461) in HeLa cells (**g**) and Hep G2 cells (**h**). **i** and **j**, Quantification of NAD(P)H relative concentration changes in different conditions. Asterisks denote significance levels that were determined by the two-sided paired *t*-test (*n* = 10 cells for each box plot). Scale bars: 30 µm.

We began by treating HeLa cells with actinomycin D (ActD), which is known to be a potent inducer of nucleolar stress via inhibition of RNA polymerase I transcription (8, 48) (Fig. 3a). After dosing HeLa cells with a low level of ActD (50 nM), we performed 25 minutes of time-lapse label-free imaging with a frame rate of 1 Hz (Fig. 2c; see Supplementary Video 1 and Methods for detailed sample preparation). From the THG signal alone, we were able to detect and quantify expected hallmarks of nucleolar stress, including increased nucleolar ellipticity and reduced nucleolar size (Fig. 2d; see Methods for calculations), consistent with what is seen in conventional phase-contrast imaging (Supplementary Fig. S6) (8, 48). Beyond these structural changes, however, we were also able to observe that different cellular compartments—the cytoplasm, nucleus, and nucleolus—exhibited distinct trends in metabolic coenzyme changes, specifically NAD(P)H and FAD levels (Fig. 2d). Due to the high resolution of our imaging and the preserved structural context, we were able to characterize metabolic changes across different time points, at the pixel level (∼400 nm) within a single cell (Fig. 2e and f).

**Fig. 3:**
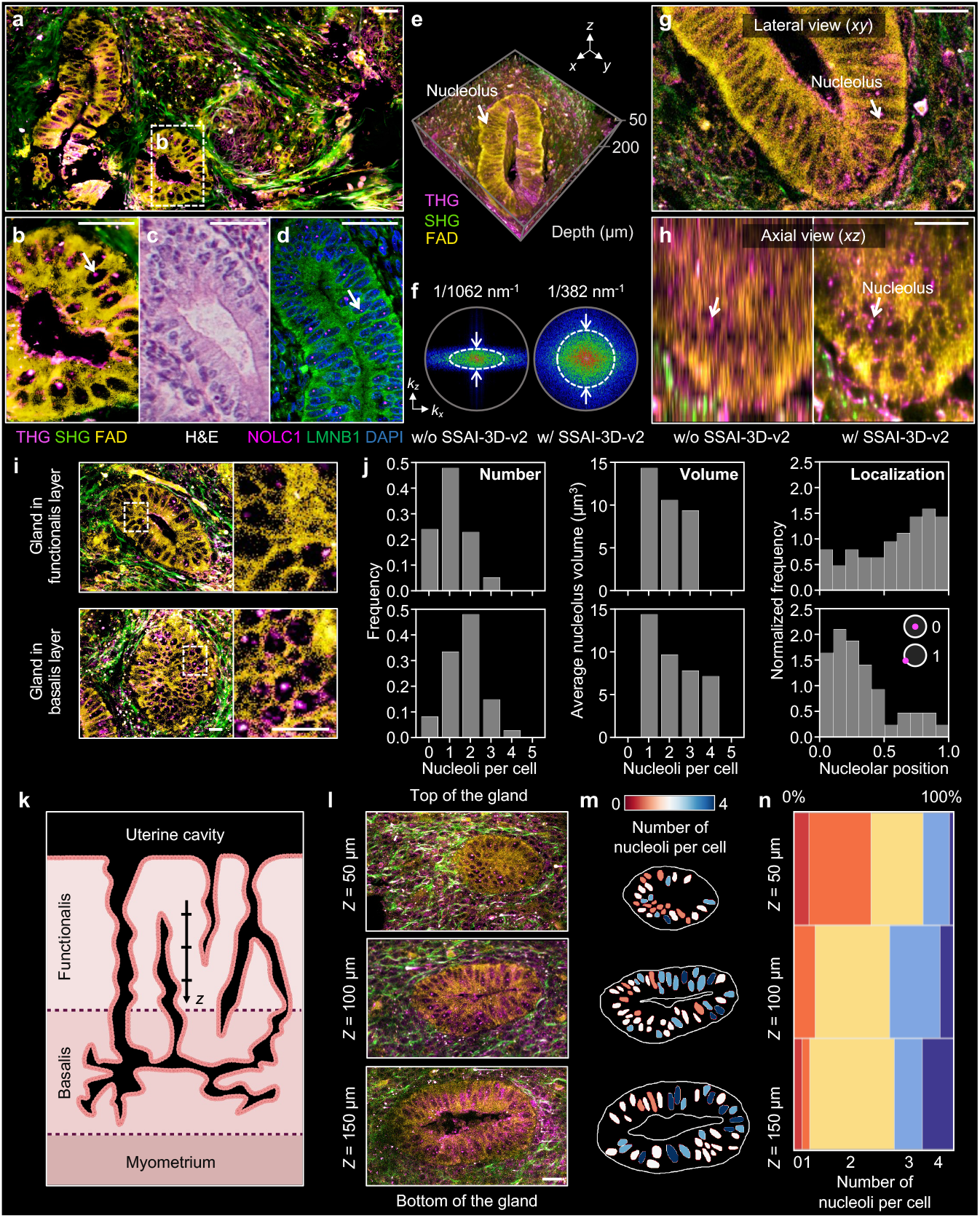
Spatial- and depth-resolved mapping of nucleolar statistics in fresh and intact human tissue. **a**, Metabolic and structural imaging of freshly excised human uterine tissue. **b-d**, Comparative images of endometrial glands from the same patient as in **a**, shown with metabolic and structural imaging (**b**), hematoxylin and eosin (H&E) staining (**c**), and immunofluorescent staining (**d**). White arrows indicate the nucleolus. **e**, 3D high-resolution structural and metabolic imaging showing an endometrial gland. **f**, Fourier spectrum of axial images before and after axial deblurring with SSAI-3D-v2, with characterized axial resolutions. **g**, Lateral image of the uterine tissue, with the nucleolus highlighted by the white arrow. **h**, Comparison of axial images of the uterine tissue with and without axial deblurring using SSAI-3D-v2, with the white arrows pointing to the same nucleolus. **i**, Representative glands from the stratum functionalis and stratum basalis in the endometrial tissue. **j**, Statistics of the glands in **i** (*n* = 50 cells). Number: the number of nucleoli per cell. Volume: the average volume of nucleoli in a cell. Localization: the distance between the nucleolus and the nuclear center of mass, normalized by the nuclear radius. A nucleolar position of 0 means the nucleolus is at the center of the nucleus, while 1 means the nucleolus is on the nuclear membrane. **k**, Schematic of the 3D human endometrium showing coiled glands extending through the stratum functionalis and stratum basalis above the myometrium. **l**, Images of the same gland at different depths, from the top (*Z* = 50 µm) to the bottom (*Z* = 150 µm) of the gland. **m**, Corresponding segmentations of the nucleus in **l**. The color of each nucleus represents the number of nucleoli in the corresponding cell. **n**, Proportions of cells with different numbers of nucleoli at various depths. Scale bars: 30 µm.

At *t* = 25 min post-ActD treatment, nucleolar NAD(P)H levels decreased by 4.3%, consistent with the cessation of nucleolar functions, while nuclear levels dropped by 16.3% and cytoplasmic levels by 0.7%. The NAD(P)H decline was not constant: at *t* = 5 min, nuclear NAD(P)H had decreased by only 1.7%, with cytoplasmic and nucleolar changes remaining below 0.5%. The pronounced nuclear decline is particularly notable, as it may reflect a broader impact of nucleolar stress, given NADH’s critical roles in chromatin decompaction, DNA replication, and DNA repair (Fig. 2e and f) (49, 50).

To further understand how nucleolar stress drives nuclear NAD(P)H depletion, we examined the role of poly(ADP-ribose) polymerases (PARPs), which are known to maintain nucleolar structure and function (51–54). PARPs relocalize from the nucleolus to the nucleoplasm in response to nucleolar stress (54–56), which could contribute to the depletion of nuclear NAD(P)H that we had observed. To test this directly, HeLa cells were treated with ActD in the presence or absence of the PARP inhibitor PJ34 (Supplementary Fig. S7). The addition of PJ34 not only caused more disordered nucleolar structure (Supplementary Fig. S7b), but was also associated with a smaller decline in nuclear NAD(P)H levels compared to the ActD-only condition (Supplementary Fig. S7c)—supporting an unexpected role for PARP in driving NAD(P)H depletion in the nuclear compartment when cells are experiencing nucleolar stress.

To assess the generalizability of our observations, we confirmed that ActD treatment also led to comparable results in HepG2 cells (Fig. 2g and h). We further validated these findings by exposing both HeLa and HepG2 cells to CX-5461, a nucleolar stress–inducing compound that inhibits RNA Polymerase I transcription through topoisomerase II poisoning, a mechanism distinct from ActD (Fig. 2g and h) (57). CX-5461 produced trends similar to those of ActD, but with a more pronounced decrease in nuclear NAD(P)H and comparatively modest changes in the cytoplasm and nucleolus (Fig. 2i and j). This enhanced nuclear depletion is consistent with CX-5461’s inhibition of DNA topoisomerase II, which can increase chromatin compaction that is also regulated by PARPs. Together, these findings show that our approach is capable of capturing a broad metabolic response to nucleolar stress across cellular compartments with high spatial and temporal resolution, with the ability to reveal compartment-specific differences that develop following different mechanisms of perturbation.

### Spatial- and depth-resolved mapping of nucleolar statistics in fresh and intact human tissue

Having validated that our platform can capture high-resolution nucleolar dynamics within simple 2D monocultures, we next shifted our focus to more complex tissues. We chose the human endometrium, the hormonally responsive and highly regenerative inner lining of the uterus, where dynamic changes in nucleolar structure have been documented and proposed to play a critical role in supporting cyclic remodeling and cellular turnover throughout the menstrual cycle (10–12). This tissue offers both biological relevance and practical accessibility for evaluating nucleolar dynamics using our imaging technology.

We started by benchmarking our label-free imaging technology against conventional hematoxylin and eosin (H&E) staining and immunofluorescent staining, using freshly excised tissue samples from a secretory phase patient undergoing a total hysterectomy (Fig. 3a-d). While H&E requires fixation and extensive tissue processing and offers limited contrasts (58), our approach delivers high-resolution 3D structural and metabolic imaging of unprocessed tissue within minutes. Notably, our method provides detailed visualization of the endometrial architecture, capturing distinct features characteristic of the secretory phase such as wide tubular glands (via metabolic imaging) and a network of collagen fibers in the extracellular matrix (via SHG) (41, 59) (Fig. 3a; see Supplementary Fig. S8 for other endometrial structures). Most importantly, THG enables precise identification of nucleoli within glandular epithelial cells (Fig. 3b), features that can be seen in immunofluorescent staining with NOLC1 (Fig. 3d) but are not readily discernible with H&E (Fig. 3c). We further validated that the observed discrete, spherical structures correspond to nucleoli through electron microscopy of the same tissue sample (Supplementary Fig. S9). Moreover, by applying our axial deblurring network, SSAI-3D-v2, we extend this high-resolution capability into three dimensions (Fig. 3e). The improved axial resolution of ∼400 nm enables accurate identification and localization of nucleoli through tissue depths exceeding 200 µm, which was not possible before axial deblurring (Fig. 3f-h).

Next, we sought to determine whether there is heterogeneity in nucleolar distribution across different endometrial glands. To investigate this, we selected two representative glands: one located in the stratum functionalis, the superficial layer of the endometrium that undergoes hormone-controlled thickening and shedding throughout the menstrual cycle; and one located in the stratum basalis, the deeper layer of the endometrium that is minimally hormone responsive and remains relatively constant (60) (Fig. 3i). For each gland, we quantified the number of nucleoli per cell, the average nucleolar volume, and the proximity of each nucleolus to the nuclear membrane—a feature previously linked to rising progesterone levels in the secretory phase (Fig. 3j; *n* = 50 cells) (61). While we observed no significant differences in nucleolar number or volume between the two glands, we found distinct statistical distributions in nucleolar positioning. More nucleoli in the functionalis gland were located closer to their respective nuclear membranes, compared to nucleoli in the basalis gland. This trend underscores that glands in the basalis layer are functionally different from the luminal epithelium, with the former serving as a reservoir for regenerating the functionalis while the latter is actively involved in progesterone-triggered secretory functions (62, 63).

We further leveraged the high axial resolution of our imaging approach to examine whether nucleolar characteristics vary within individual epithelial cells of a single gland. We selected an occluded gland from the stratum functionalis, an architecture that is found in greater abundance in the secretory phase compared to the proliferative phase (Fig. 3k and l) (60, 64). Depth-resolved 3D mapping over 150 µm revealed that epithelial cells located near the base of the gland exhibited fewer nucleoli than those positioned closer to the endometrial lumen (Fig. 3m and n). This spatial variation is consistent with known endometrial biology, where luminal and upper glandular epithelial cells exhibit higher proliferative activity, particularly during the estrogen-driven proliferative phase of the menstrual cycle (65, 66). Taken together, these findings demonstrate that our label-free structural and metabolic imaging platform not only enables 3D visualization of global tissue architecture, but also uncovers rich, quantitative subcellular information—revealing biologically relevant features at an unprecedented resolution that are inaccessible by conventional methods.

### Context-dependent nucleolar translocation and cellular metabolic adaptations to hormone signaling in endometrial organoids

Building on the nucleolar studies that we performed in tissues, we next asked if our imaging system could be used to explore how these spatial nucleolar distributions in the endometrium form over time. Using a tissue-engineered model of the endometrium, we sought to investigate how physiological hormonal perturbations can modulate nucleolar organization and cell states, with the goal of identifying dynamic structural patterns and metabolic signatures that may yield diagnostic insights. To accomplish this, we adapted our previously published protocol for 3D synthetic hydrogel co-culture of primary, donor-matched endometrial epithelial organoids and endometrial stromal cells—an *in vitro* model shown to recapitulate key morphological and functional changes occurring throughout the human menstrual cycle (Fig. 4a and b) (67).

**Fig. 4:**
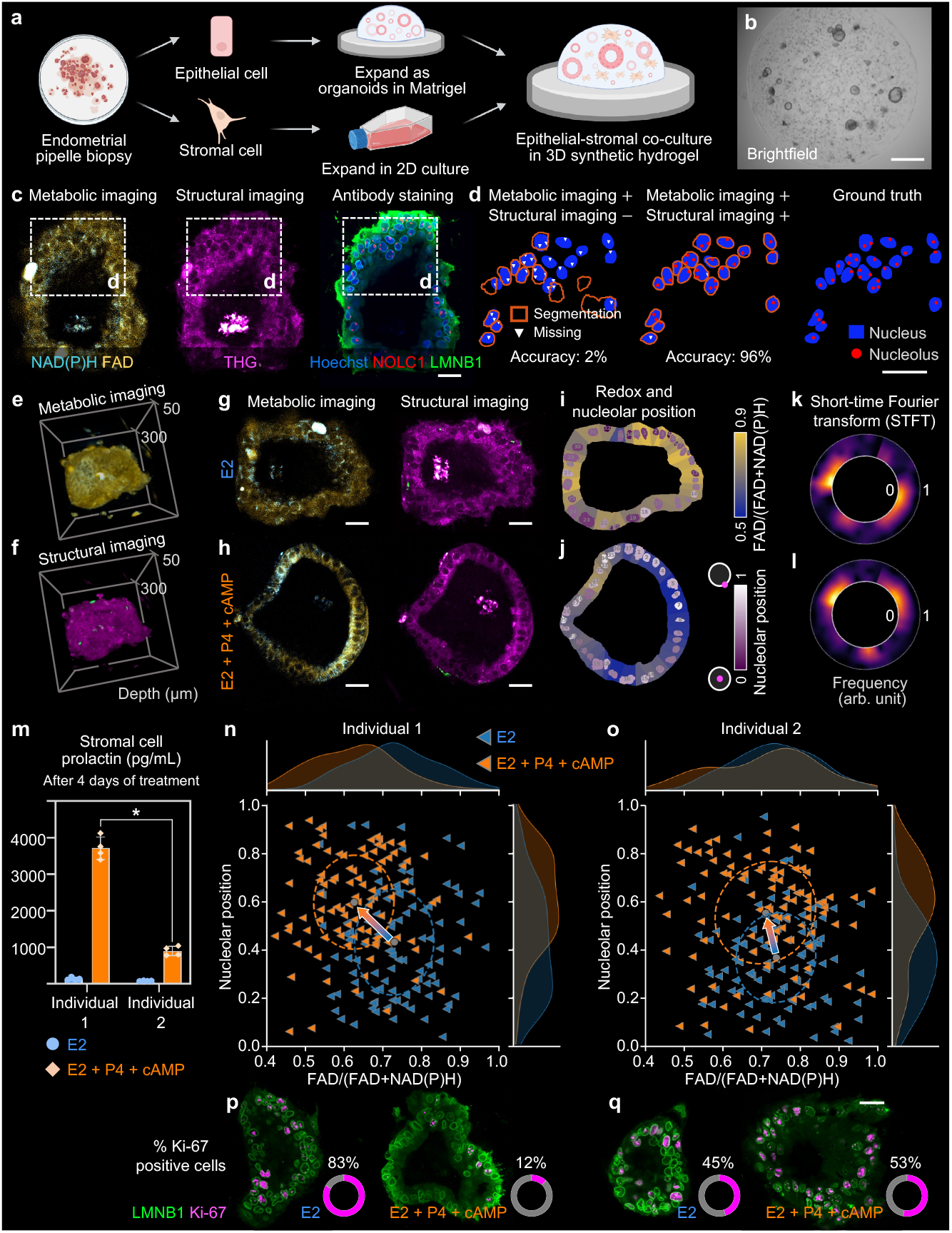
Context-dependent nucleolar translocation and cellular metabolic adaptations to hormone signaling in endometrial organoids. **a**, Endometrial epithelial organoids and stromal cells isolated from pipelle biopsies were expanded and encapsulated within a 3D synthetic hydrogel. **b**, Representative brightfield image of the setup. **c**, Co-registered images of metabolic and structural imaging, as well as antibody staining (fixed and stained after live label-free imaging) of an endometrial organoid. **d**, Validation of nucleus segmentation and nucleolus detection. Right: ground truth derived from antibody staining. **e**-**f**, Example of 3D metabolic and structural imaging of an endometrial organoid. **g**–**h**, Representative metabolic and structural images from the control (E2) and treated (E2 + P4 + cAMP) conditions to mimic the proliferative and secretory phases of the menstrual cycle, respectively. **i**–**j**, Quantifications of images in **g** and **h**, where the color of the cytoplasm represents the optical redox ratio, and the color of the nucleus represents the nucleolar position. **k**–**l**, Short-time Fourier transform (STFT) analysis of redox ratios. **m**, Measurements of prolactin secreted by stromal cells derived from two individuals after 4 days with different treatment conditions in hydrogel mono-culture. **n**–**o**, Nucleolar position versus optical redox ratio in endometrial epithelial organoid cells from Individuals 1 and 2 under the two different treatment conditions. The ellipses in the plots represent 2D Gaussian fits, and the distributions outside the plots show the 1D distributions in two axes. **p**–**q**, Antibody staining and quantified percentage of Ki67-positive cells from individuals 1 and 2 under different treatment conditions. Scale bars: 0.5 mm (**b**); 30 µm (**c, d, g, h, p, q**).

We first demonstrated that live organoids embedded in hydrogel could be imaged with high fidelity using our multimodal imaging approach, which enabled the detection of cellular metabolism via NAD(P)H and FAD autofluorescence, as well as nucleolar structures through THG signals (Fig. 4c). To this end, we cross-referenced the same organoids following immunofluorescence staining with antibodies against NOLC1. Given that nucleolar dynamics in the endometrium often involve changes in nucleolar positioning within the nucleus, we further stained the nuclear lamina using an antibody against LAMINB1 (LMNB1), a structural component of the nuclear envelope (Fig. 4d; see Supplementary Fig. S10 for pipeline). Not only were we able to identify nucleoli with 96% accuracy, but we also resolved their spatial localization within the nucleus (Fig. 4d; see Supplementary Fig. S11 for a 3D example). These results establish that our label-free imaging platform can effectively resolve nucleolar architecture and positioning within endometrial organoids that are cultured alongside stroma in a complex, 3D microenvironment.

We next turned to investigating whether our imaging method is sensitive enough to capture dynamic changes in endometrial organoids in response to progesterone treatment (Fig. 4e-l). Progesterone plays a critical physiological role in preparing the endometrium for potential embryo implantation; during the secretory phase of the menstrual cycle, a rise in progesterone is associated with reduced epithelial proliferation and alterations in nucleolar structure and localization (Supplementary Fig. S12a) (68, 69). To mimic the secretory phase, the epithelial organoid-stromal cocultures were treated with a combination of estradiol (E2), progesterone (P4), and cyclic adenosine monophosphate (cAMP) (Fig. 4h), while organoids exposed to E2 alone served as a control to simulate the proliferative phase (Fig. 4g) (70–72).

We used measurements of NAD(P)H and FAD autofluorescence to calculate the optical redox ratio (FAD/(FAD+NAD(P)H)), which has been shown to correlate with metabolic activity and, by extension, may reflect cellular proliferation (see Supplementary Fig. S13 and Methods for calculation) (36–40). Based on this analysis, control organoids (treated with E2 alone) remained highly metabolically active (Fig. 4i), whereas organoids exposed to progesterone exhibited a decreased optical redox ratio (Fig. 4j)—likely reflecting reduced proliferation, consistent with known *in vivo* effects of progesterone. These observations align with our previous bulk transcriptomic data from epithelial-stromal co-cultures, where Gene Ontology (GO) enrichment analysis identified “FAD binding” and “NADH dehydrogenase activity” as processes significantly upregulated and downregulated, respectively, in response to progesterone treatment (Supplementary Fig. S11) (67). Interestingly, cellular metabolic activity within the control organoids was not uniformly distributed. We observed striking spatial asymmetry in redox potential, with discrete yet contiguous regions displaying either high or low metabolic activity (Fig. 4k and l). This metabolic heterogeneity was strongly correlated with nucleolar positioning within the nucleus: in regions of elevated redox activity (higher optical redox ratio), nucleoli were more frequently centrally localized. In contrast, in cells with lower redox activity (lower optical redox ratio), nucleoli shifted toward the nuclear periphery, presumably closer to the nuclear envelope. In progesterone-treated organoids, this redox asymmetry became more spatially restricted, and nucleoli predominantly localized to the nuclear periphery (Fig. 4i and j). We further confirmed, at the ultrastructural level using CLEM, that nucleoli are more likely to localize closer to the nuclear envelope in response to progesterone treatment (Supplementary Fig. S14). Together, these findings demonstrate that progesterone modulates both the metabolic state and nucleolar architecture of endometrial organoids. More broadly, they underscore the power of our label-free imaging platform to spatially resolve metabolic heterogeneity and link these profiles to dynamic changes in nucleolar localization at single-cell resolution within complex 3D tissue models.

Next, we sought to explore the potential diagnostic utility of our imaging technology by examining differences in the relationship between nucleolar dynamics and cellular metabolism in endometrial organoids derived from two distinct tissue donors. The first donor (Individual 1) presented with a small intramural fibroid, while the second donor (Individual 2) was diagnosed with Stage IV endometriosis, characterized by endometriotic tissue infiltrating the uterine serosa and ovaries and severe fibrovascular adhesions tethering the uterus to the sigmoid colon and upper rectum. For this comparison study, we first sought to differentiate between the donors by measuring stromal cell prolactin secretion, a key downstream marker of progesterone signaling and a critical mediator of endometrial function. In hydrogel mono-cultures, we observed that stromal cells derived from Individual 2 were significantly less responsive to progesterone, recapitulating the aberrant response to progesterone—termed “progesterone resistance”—often described in endometriosis patients (Fig. 4m).

Given that stromal-epithelial crosstalk is a key mediator of progesterone’s effects on epithelial cells (67), we next examined organoids derived from these two individuals within our hydrogel co-culture setup. Upon progesterone treatment, endometrial organoids from Individual 1 exhibited a marked decrease in optical redox ratio—indicative of metabolic reprogramming consistent with the cessation of proliferation (Fig. 4n). In contrast, organoids from Individual 2 failed to show a comparable metabolic shift, suggesting an impaired ability to respond to progesterone-mediated signals (Fig. 4o). Importantly, although organoids from both donors exhibited dynamic changes in nucleolar positioning, only organoids from Individual 1 showed consistent and uniform directional shifts in nucleolar localization (Fig. 4n). In contrast, organoids from Individual 2 displayed a more heterogeneous and dispersed pattern, further indicating a blunted or dysregulated response to progesterone (Fig. 4o). These findings suggest that the combined analysis of redox state and nucleolar repositioning provides a sensitive and functional readout of progesterone responsiveness in endometrial organoids co-cultured with donor-matched stroma.

To directly assess the proliferative response to progesterone, we evaluated Ki67 expression—a well-established marker of cell proliferation—in organoids derived from both individuals. Consistent with predictions based on our label-free metabolic imaging, progesterone treatment significantly suppressed epithelial proliferation in co-culture organoids from Individual 1, reducing the proportion of Ki67-positive cells from 83% to just 12% (Fig. 4p). In contrast, organoids from Individual 2 showed a blunted response, with 53% of cells remaining Ki67-positive following treatment (Fig. 4q). This outcome is consistent with impaired progesterone responsiveness and mirrors clinical features associated with progesterone resistance.

Taken together, these findings highlight the diagnostic potential of our labelfree imaging platform to detect nuanced, yet biologically meaningful, differences in hormone responsiveness across patient-derived organoids—enabling high-resolution, spatially resolved assessments of metabolic state and nucleolar dynamics within complex, 3D tissue-engineered models.

## Discussion

The multiscale capability of our label-free imaging platform—spanning single cells, patient-derived organoids, and intact tissues—provides critical insights into nucleolar dynamics and cell state responses to physiologically relevant perturbations, particularly within the 3D context of hormonally responsive endometrial organoids and tissue. This approach supports a long-standing hypothesis in the field: that the nucleolus, due to its essential role in cell growth and proliferation, can serve as a diagnostic marker in disease (73–75). Our study provides a proof of concept that, through this non-invasive imaging technology, this potential can now be translated into practical application.

The human endometrium presents a highly relevant context for studying nucleolar biology, as dynamic changes in nucleolar structure and localization are closely linked to fluctuations in the steroid hormone progesterone, which plays a critical role during the secretory phase in regulating endometrial regeneration and preparation for embryo implantation. Dysregulated progesterone signaling, often in the form of “progesterone resistance,” has been strongly linked to impaired decidualization of endometrial stromal cells, unchecked cellular proliferation, and persistent inflammation, all of which contribute to the pathogenesis of several endometrial disorders including endometriosis, adenomyosis, endometrial hyperplasia, and implantation failure (68, 69, 76). However, the biological significance of progesterone-induced nucleolar remodeling in the endometrium—and the potential of the nucleolus to serve as a biomarker in this context—has remained largely unexplored. By allowing precise visualization of nucleoli, quantification of their spatial localization, and integration of this information with proliferative status across different tissue depths, we were able to dissect the effects of progesterone on nucleolar behavior and tissue metabolism. We further extended our analysis to organoids derived from an individual exhibiting stromal progesterone resistance, showing that our noninvasive imaging approach successfully captured nucleolar and metabolic alterations associated with progesterone insensitivity at both the cellular and subcellular levels.

Altogether, this study highlights the power of our imaging platform in advancing the study of nucleolar dynamics in tissues. By bridging the gap between simplified cell culture models and complex, physiologically relevant 3D tissues, our approach facilitates the integration of structural, functional, and metabolic information. This enables a more holistic understanding of cell states in health and disease and opens new avenues for identifying diagnostic or therapeutic biomarkers in hormone-responsive tissues. While further validation in tissues and organoid systems beyond the endometrium is needed, these early findings suggest that label-free structural and metabolic imaging can complement traditional biomarkers, particularly in contexts where labeling is limited or ambiguous (59, 77). This advancement positions imaging as an active tool for identifying, quantifying, and manipulating cellular features, ultimately enhancing diagnostic and therapeutic strategies.

## Methods

### Multimode fiber source

The parameters of the multimode fiber source were further optimized for the imaging system, building upon our previous design (Supplementary Fig. S1) (34). A step-index multimode fiber (Thorlabs, FG025LJA; 25 µm core diameter, 0.10 NA) of 7.4 cm length was launched by a femtosecond mode-locked ytterbium laser (Light Conversion, Carbide). The pump laser operated at 1030 nm with a pulse width of 219 fs, where 240 nJ of pulse energy was used for focusing. To match the 18.2 µm mode field diameter of the fiber, an achromatic doublet with a 60 mm focal length (Thorlabs, AC254-060-B) was employed for fiber coupling, achieving a coupling efficiency of 90%. The multimodal nonlinear pulse propagation was controlled by a fiber shaper device (34, 45), which was driven by a single stepper motor-based actuator (Hilitand, 2-Phase 4-Wire Stepper Motor) with a microcontroller unit (Arduino, Uno Rev3). The motion resolution was 25 µm. To collimate the output beam and band-select the light centered at 1100 nm for imaging, the output supercontinuum was collimated by an off-axis parabolic mirror with a 76 mm focal length (Edmund Optics, 36-588) and spectrally filtered by a 1100±25 nm bandpass filter (Edmund Optics, 85-906).

### Structural and metabolic imaging

The structural and metabolic imaging was implemented using an inverted scanning microscope (Supplementary Fig. S1) (34). A continuously variable neutral density filter (Thorlabs, NDC-50C-4M-B) was placed after the multimode fiber source and before the imaging system to adjust the pulse energy delivered to the sample. The beam was scanned by a galvanometer mirror pair (ScannerMAX, Saturn) and conjugated to the back focal plane of the objective (Olympus, XLPLN25XWMP2). The emission path was separated from the excitation path by a dichroic mirror (Thorlabs, DMLP650L) and was further divided into four detection channels using dichroic mirrors (Chroma, T412lpxt, T505lpxr, T570lpxr) and bandpass filters (Chroma, ZET365/20x; Edmund Optics, 84-095; Edmund Optics, 65-159; Semrock, FF01-609/57-25), corresponding to THG, NAD(P)H, SHG, and FAD. The emissions were collected by four individual photomultiplier tubes (Hamamatsu, H16201) and amplified by transimpedance amplifiers (Femto, DHPCA-100). The signals were converted into images using custom-written acquisition software. For 3D imaging, the software automated data acquisition with a microscope stage (ASI, MS2000) that adjusted the sample positions. The pulse energy at the excitation focus was maintained at a constant level at different depths by adjusting the neutral density filter to compensate for light attenuation in the samples. All image visualizations were generated in the Fiji software, with the 3D viewer used for 3D volumes, and magenta (hot), cyan (hot), green, and yellow (hot) rendered for the THG, NAD(P)H, SHG, and FAD images, respectively.

### Optical redox ratio calculation

Prior to the calculation of the optical redox ratio, fluorescence intensities were normalized by considering the multiphoton generation efficiency and the detector gain. Standard solutions of FAD (MilliporeSigma, F6625) and NAD(P)H (MilliporeSigma, N8129) were prepared (1 mM each) to validate the multiphoton origin of the signals (Supplementary Fig. S2) and to calibrate the optical redox calculation (78). Following conventions from previous literature, the optical redox ratio was defined as the ratio of the concentrations of FAD and NAD(P)H (34, 41, 79)

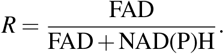

To calculate the redox ratio for a specific cell or a particular compartment within a cell, automated segmentation was first performed to generate a binary mask (as described below). The concentrations of FAD and NAD(P)H were then determined by summing the pixel values within the binary mask. In Fig. 2d, to better visualize the concentration trend and reduce noise contamination, the resulting curve was smoothed using a rolling average with a window length of 3. In Fig. 2e, to better visualize the rate of concentration change, the raw heatmap was convolved with a 2×2 kernel of ones.

### Optical characterizations

The supercontinuum spectrum was measured using a NIR spectrometer (Ocean Insight, NIRQuest+1.7). The near-field output spatial profile of the multimode fiber was measured using a complementary metal-oxide semiconductor (CMOS)-based camera (Mako, G-040B) positioned after a 4-F relay system (80). The spatial profiles at different bands were measured with corresponding spectral filters. The 3D resolution of the imaging system was characterized by imaging 0.1 µm green and 0.2 µm blue sub-diffraction-limited fluorescent microspheres (BangsLabs, FCDG002 and FCGB003), which were fitted with a 2D Gaussian distribution in both the lateral and axial planes. The full width at half maximum (FWHM) of the fitted distribution was used to represent the resolution. The comparison of 3D resolution before and after axial deblurring was performed using the Fourier transform spectrum of the axial image.

### Axial deblurring with SSAI-3D-v2

Axial deblurring with SSAI-3D-v2 was inspired by our previous framework, SSAI-3D (42), while focusing on further improving reconstruction fidelity by increasing the training data size and minimizing domain shift with more diverse cellular data. SSAI-3D-v2 employed a self-supervision strategy and enabled instant inference of axial deblurring without training with the single-stack data. The training of SSAI-3D-v2 was conducted in two steps. First, a self-supervised paired dataset was generated by applying Gaussian blurring to a ∼10 GB cellular dataset using a set of 25 PSFs, which combined five different sizes and five different orientations. The cellular dataset was a collection of historical data from the system, and it could be replaced with any other dataset for customization. Similar to SSAI-3D, an optional denoising step was employed to enhance the quality of the training data. Second, the pre-trained deblurring network NAFNet (81) was fine-tuned using the generated self-supervised paired dataset. The sparse fine-tuning strategy was applied with the previously released pre-trained surgeon network (42). With the weights of the axial deblurring network adjusted through these two steps, a newly generated 3D single stack could be axially deblurred by sequentially inputting the axial images. The deblurred axial images in both the *xz* and *yz* directions were combined by averaging to generate the final volume. To investigate and compare the performance of overcoming domain shift, simulations with very low lateral-axial similarity were performed in Fig. 2d by generating cylinders with random sizes, locations, and intensities within a 224 µm × 224 µm × 224 µm volume.

### Network training and evaluations

To adapt the microscopy images to the pre-trained network, which required natural scene RGB input, multi-channel microscopy images were separated into several single-modality grayscale images, and each channel was replicated to create an RGB image that matched the format of the pre-trained network. The RGB predictions were then inversely averaged along the color dimension to convert them back into a single-modality grayscale image. The network took input patches of 256 × 256 with a batch size of 8. For all presented experiments, network parameters were optimized using the Adam optimizer with a learning rate of 10^−3^ and a weight decay of 10^−4^. Training was performed on a single Nvidia GeForce RTX 3080 card, and the model was quantized to float16 precision for memory efficiency. With the sparse fine-tuning strategy, approximately 90% of the layers in the pre-trained network were frozen, while only about 10% were fine-tuned.

We compared the performance of SSAI-3D-v2 with three baseline methods: Self-Net (82), CARE (83), and SSAI-3D (42). All methods were implemented using the open-source codes released with their respective papers. Since CARE was the only method among them that required a known PSF model, a Gaussian blur with a standard deviation of 5 was assumed for the CARE framework (83). For all presented experiments, the hyper-parameters and training settings remained consistent. We used MSE to compare reconstruction fidelity.

### Image segmentation, co-registration, and analysis

Image segmentation was performed using the recent foundational segmentation model SAM (Segment Anything Model), which was pre-trained with 11M images (SA-1B) (84). To increase the specificity of segmentation, manually labeled bounding boxes were provided as prompts to the segmentation model. To enable 3D analysis, the labeled masks at each layer were further indexed and aligned across different layers, resulting in the 3D segmented masks.

Structural and metabolic images were inherently co-registered due to the use of a single excitation. The co-registration of images across different systems was achieved in four steps. First, a bright-field image was taken to roughly locate the regions of interest. Second, label-free structural and metabolic imaging was performed on the cells in their living states. Third, samples were fixed and stained with the desired dyes and antibodies (see below for more details on fixation and staining protocols). They were then imaged using confocal microscopy, referenced against the initial bright-field image for rough alignment with the label-free images. Lastly, with the recorded depth from both systems, images were aligned using customized software to achieve pixel-wise co-registration, accounting for rotation and differences in pixel sizes between the two systems.

Using the 3D segmented mask, the center of the volume was determined as the center of mass. The localization of the nucleolus was then calculated based on its distance from the center and normalized by the radius, assuming a spherical shape. The size of the nucleolus in the unit of length was estimated by fitting a 1D Gaussian function to the line profile, while its volume was estimated using a 3D Gaussian fitting. In Fig. 2d, ellipticity was quantified as the ratio of the short axis to the long axis using a 2D Gaussian fit. In Fig. 4k and l, the STFT analysis was performed on the ordered redox data with a window size of 3. In Fig. 4p and q, under the same imaging settings, Ki-67 intensities were thresholded by a value to mark them as Ki-67 positive or negative.

### 2D cell culture and experiments

The methods for culturing and treating HeLa and HepG2 cells were primarily adopted from a previous publication (54). HeLa S3 cells (ATCC) and HepG2 cells (ATCC) were cultured under standard conditions (37°C, 5% CO2) in high-glucose DMEM (GenClone, 25-500) supplemented with 10% FBS (Gemini Bio) and 1X Pen-Strep-Glutamine (Life Technologies, 10378016). For nucleolar perturbation experiments, actinomycin D (Sigma, A1410-5MG) and CX-5461 (Selleckchem, S2684) treatment were administered by directly adding the compound to the culture medium at a concentration of 50 nM and 30 µM, respectively, for 30–60 minutes. The chemicals were diluted in high-grade DMSO Hybrid-Max (Millipore-Sigma, D2650). For experiments involving PARP inhibition (Supplementary Fig. S7), PJ34 (20 µM; Selleckchem, S7300) was added 16 hours prior to ActD treatment. For experiments to validate NAD(P)H autofluorescence signal, HeLa cells were treated with 10 nM daporinad (Selleckchem, S2799) for 30 minutes.

### Endometrial tissue acquisition and cell culture

All endometrial tissue samples were collected from participants who provided informed consent under a protocol approved by the Partners Human Research Committee and the Massachusetts Institute of Technology Committee on the Use of Humans as Experimental Subjects (Protocol number IRB-P001994). For label-free imaging of intact human endometrium, freshly excised uterine tissue was obtained from a 45-year-old patient undergoing a total hysterectomy, with histological confirmation of benign secretory endometrium. After imaging, tissue sections were fixed in 4% paraformaldehyde (PFA) and stored after imaging (85).

The methods for culturing primary human endometrial cells were adapted from our previous study (67). Briefly, endometrial tissue was collected via pipelle biopsies from reproductive-age women undergoing laparoscopic surgery for non-malignant gynecologic conditions. Endometrial epithelial glands and stromal cells were isolated from biopsies through enzymatic digestion and cultured separately, either as endometrial epithelial organoids (EEOs) in 25 µL Matrigel droplets or as endometrial stromal cells (ESCs) in traditional cell culture flasks. EEOs were cultured in our previously published EEO expansion medium containing advanced basal media cocktail (67) (ABM, 1X); rhEGF (50 ng/mL, Corning); rhNoggin (100 ng/mL, Peprotech); rhRspondin-1 (200 ng/mL, R&D systems); rhFGF-10 (50 ng/mL, Peprotech); Nicotinamide (1 mM); Insulin-Transferrin-Selenium (ITS, 1%, Invitrogen); N-acetyl-L-cysteine (1.25 mM); TGFB pathway inhibitor (A83-01, 500 nM, Peprotech); Rock inhibitor Y-27632, (ROCKi, 10 mM, Tocris), 17*β* -estradiol (E2, Sigma Aldrich) E2 (1 nM). ESCs were cultured in phenol red-free DMEM/F12 with 5% charcoal-stripped calf serum, 1 nM E2, and 1x Pen-Strep solution (Sigma Aldrich). For this study, cells were isolated from two donors: a 47-year-old patient with a small intramural fibroid (“Individual 1”), and a 45-year-old patient with Stage IV endometriosis, characterized by endometriotic tissue infiltrating the uterine serosa and ovaries and severe fibrovascular adhesions tethering the uterus to the sigmoid colon and upper rectum (“Individual 2”).

### Endometrial epithelial-stromal co-culture

Epithelial-stromal co-culture experiments were set up following previously published protocols unless otherwise noted (67). Briefly, donor-matched intact EEOs (6 days after passaging from single cells, ∼100 µm in diameter) and ESCs were harvested via Matrigel dissolution and trypsinization, respectively. After counting, EEOs and ESCs were combined into a single cell pellet and resuspended in a synthetic hydrogel precursor solution comprising 8-arm polyethylene glycol-vinyl sulfone (PEG-VS) macromers functionalized with peptides that bind endometrium-relevant integrins and cell-secreted extracellular matrix. A matrix metalloproteinase-degradable crosslinker was added to initiate gelation. After gentle mixing, the cell-laden hydrogel solution was pipetted in 3 µL droplets in glass-bottom 96-well plates (CellVis, P96-1.5H-N). The droplets, which had a final count of 30 EEOs and 10,000 ESCs per 3 µL droplet, were incubated for 30 min in a humidified incubator at 37°C until gelation was complete.

The droplets were cultured for 2 days in a 1nM E2 modified EEO medium formulation, with the Rock inhibitor Y-27632 and TGF-*β* inhibitor A83-01 omitted from the EEO expansion medium formulation due to the co-culture format with intact organoids in synthetic hydrogel. After 2 days, the wells were changed to modified EEO medium containing either 10nM E2 (“E”) to mimic the proliferative phase of the menstrual cycle, or 10nM E2, 1 µM progesterone (P4, Sigma-Aldrich, 5341), and 500µM 8-bromoadenosine 3’,5’-cyclic monophosphate (cAMP, Sigma-Aldrich, B7880) (“EPc”) to mimic the secretory phase. After 4 days of E or EPc treatment (with medium refreshed on day 2), the co-cultures were imaged and then fixed with 4% PFA for later immunofluorescent staining. ESC mono-cultures from the two biopsy donors were also set up with the same hydrogel formulation and ESC concentration to characterize responsiveness to EPc treatment. Cell culture supernatant was collected every 2 days; prolactin levels were measured from supernatant collected on day 4 using a prolactin ELISA DuoSet (R&D Systems/Fisher, DY682).

### Immunofluorescent staining and confocal imaging

HeLa cells were fixed directly in the culture medium with 4% PFA for 15 minutes at room temperature, followed by multiple PBS washes and storage at 4°C. For organoid fixation, samples were incubated with 4% PFA for 30 minutes at room temperature on an orbital shaker, followed by multiple PBS washes and storage at 4°C. For staining of endometrial tissue, hysterectomy specimens were fixed for at least 24 hours in 4% PFA before being embedded in paraffin blocks. Paraffin-embedded tissue sections were cut, mounted on slides, and deparaffinized using the automated slide processor. Slides were rinsed twice with PBS and incubated in PBS for an additional 5 minutes. Antigen Retrieval: Slides were rinsed in antigen retrieval buffer (10 mM sodium citrate, pH 6.0; prepared by mixing 1.8 mL of 100 mM citric acid and 8.5 mL of 100 mM sodium citrate, and bringing to a final volume of 100 mL with distilled water). Fresh antigen retrieval buffer was added to a Coplin jar containing the slides, which was loosely covered and placed in a heat sink. Antigen retrieval was performed in a 1250 W microwave at 60% power for 6 minutes, followed by three 3-minute intervals at 40% power. Buffer was replenished between intervals to compensate for evaporation. Slides were allowed to cool for 25 minutes at room temperature and rinsed in PBS.

During staining, samples were first blocked overnight in PBSA buffer (PBS containing 1% BSA, 0.1% Triton X-100, and 0.05% sodium azide). The primary antibodies used in this study included NOLC1 (Santa Cruz Bio, sc-374033), LAMIN B1 (Abcam, ab16048), and PE-Ki-67 (Miltenyi Biotec, REA183), each diluted 1:200 in PBSA. The corresponding secondary antibodies, Alexa Fluor 647 (ThermoFisher, A-27040) and Alexa Fluor 488 (ThermoFisher, A-11008), were diluted 1:1000 in PBSA (1x PBS, 0.1% Triton-X). Nuclei were stained with a 1:500 dilution of Hoechst (ThermoFisher, H1399) or 300nM DAPI (ThermoFisher, D1306).

Confocal imaging was performed using a Zeiss LSM 880 confocal laser scanning microscope. Images of different fluorophores were acquired sequentially using their respective excitation wavelengths for optimal performance. To facilitate alignment with label-free images, a 63X objective was used with a sampling pixel size of ∼200 nm.

### Correlative light and electron microscopy (CLEM)

Cells were fixed by adding 4% PFA directly to the culture wells at a 1:1 ratio with the existing culture medium, resulting in a final concentration of 2% PFA. Fixation was performed at room temperature for 30 minutes. The PFA/media mixture was then removed, and fresh 4% PFA in buffer was added to the wells to ensure that samples did not dry out. Cells were fixed overnight at 4°C.

Hydrogel co-culture samples were fixed by diluting an equal volume of culture media with 4% paraformaldehyde (PFA) (Electron Microscopy Sciences, EMS 15700) in 0.1M PBS buffer, resulting in a final PFA concentration of 2%. This mild fixation was carried out for 30 minutes at room temperature. Subsequently, the solution was replaced with 4% PFA in 0.1M PBS and incubated for 24 hours at 4°C. Following fixation, samples were washed three times in 0.1M piperazine-N,N’bis(2-ethanesulfonic acid) (PIPES) buffer for 5 minutes each and then quenched in 0.15% glycine (Sigma, G8790) for 10 minutes at room temperature. Samples were infiltrated with 12% pre-warmed gelatin (Sigma, G2500) for 30 minutes at 37°C. The gelatin-embedded samples were then solidified on ice for another 30 minutes before cutting it into small blocks (∼0.5 mm^3^), which were subsequently infiltrated with 2.3M sucrose (Sigma, S0389) overnight at 4°C under gentle shaking. The infiltrated blocks were then mounted onto cryo-ultramicrotomy pins (EMS, 75959-06) and frozen in liquid nitrogen where they were stored until sectioning. Sectioning was performed using a Leica UC7 cryo-ultramicrotome equipped with a cryo-diamond knife (DiATOME, cryo-immuno 35°). Semi-thin sections (1 µm) were cut at −80°C and collected onto 12mm glass coverslips using a mixture of equal parts of 2.3M sucrose and 2% methylcellulose (Sigma, M6385-100G). Sections were stored at room temperature until further processing.

For immunofluorescence, coverslips with cryosections were incubated in 0.1 M PIPES at 37°C for 30 minutes to remove the sucrose–methylcellulose layer. Samples were then blocked in 1% BSA-c (Aurion, 900.099) in 0.1 M PIPES for 30 minutes at room temperature before being incubated with primary antibodies overnight at 4°. After blocking again in the same solution, the samples were incubated with fluorescently labeled secondary antibodies for 1 hour at room temperature. Finally, coverslips were washed three times in 0.1M PIPES, followed by a final wash in deionized water, and mounted on glass slides using DAPI-containing mounting medium (Invitrogen, D3571) for nuclear staining. The samples were imaged using a Zeiss LSM 880 confocal microscope. The primary and secondary antibodies used for CLEM were identical to the ones listed above for standard immunofluorescent staining, with the addition of fibrillarin (Millipore-Sigma Anti-Fibrillarin Antibody, clone 38F3), which is 1:200 diluted in PBSA.

After fluorescent imaging, the slides were unmounted and stained with heavy metal for electron microscopy. Briefly, the samples were exposed to reduced osmium (2% osmium tetroxide, 2.5% potassium ferrocyanide) for 15 minutes, washed three times with water, and then stained again for 15 minutes with a solution of 1% TCH (thiocarbohydrazide, Sigma 223220) in water. The samples were then stained in an aqueous solution of 2% osmium tetroxide on ice, washed three times in water, stained for 15 minutes in 1% uranyl acetate in 50mM maleate buffer, washed again three times in water before undergoing serial dehydration in increasing ethanol concentrations, and resin infiltration. The infiltrated sample were baked for at least 48 hours at 60°C before being trimmed and sectioned with a Leica UC7 ultramicrotome. A diamond knife (DiATOME) was used to cut ultrathin sections (60 nm), which were then collected in carbon coated formvar slot grid (EMS, FCF2010-Cu-EA), and imaged in a Technai T12. The electron microscopy and light microscopy images were correlated using the Fiji plugin BigWarp (86).

Electron microscopy images for the HeLa cells and endometrial tissue were acquired according to the same CLEM protocol, following the same steps for cryo-ultramicrotomy.

### Statistics and reproducibility

All box plots adhere to the standard format: box bounds represent the upper and lower quartiles, lines within boxes indicate medians, and whiskers extend to data points within 1.5 times the IQR, with outliers plotted individually beyond this range. Network training and inference was independently repeated at least 3 times under different random seeds, all achieving similar results. In Fig. 2, significance were determined by the two-sided paired *t*-test. *P* values below 0.05 were considered statistically significant.

## Data availability

The data from human tissue is available at https://zenodo.org/records/13952549. Other datasets are available from the corresponding author upon reasonable request.

## Code availability

The implementations of SSAI-3D-v2 as well as some examples will be publicly available at https://github.com/You-Lab-MIT/SSAI-3D.

## Acknowledgements

The work was supported by MIT startup funds, Novo Nordisk Research Development US, Inc., NSF CAREER Award (2339338), CZI Dynamic Imaging via Chan Zuckerberg Donor Advised Fund (DAF) through the Silicon Valley Community Foundation (SVCF), the National Institutes of Health (NIH U01 EB029132 and R01 HD110335), the John and Karine Begg Fund, the Manton Foundation, R35GM142634 (E.C.), and P30-CA14051 (E.C.). We thank Tong Qiu, Matthew Yeung, Steven F. Nagle, and Haidong Feng for helpful discussions in imaging. We would also like to thank the Koch Institute Frontier Research Program, the Casey and Family Foundation Cancer Research Fund, the Koch Institute Stem Cell Initiative and Fondation MIT, the Microscopy Facility at the Swanson Biotechnology Center, the Peterson (1957) Nanotechnology Materials Core Facility, and the Hope Babette Tang (1983) Histology Facility. We are also grateful for our clinical collaborators from the Newton-Wellesley Hospital (NWH) Center for Minimally Invasive Gynecologic Surgery (Dr. Keith Isaacson, Dr. Peter Movilla, and Mollie O’Brien) for constructive discussions and facilitating access to clinical samples, as well as Maria F. Hernandez for her assistance in donor characterization of endometrial cells. Finally, we thank the NWH patients who generously consented to donate tissue samples in support of this research.

We acknowledge that the figures contain materials from BioRender (https://biorender.com/). K.L. and H.C. acknowledge support from the MathWorks Fellowship. K.L. acknowledges support from the MIT Health and Life Sciences Collaborative (HEALS) Fellowship. E.L.K. acknowledges support from the National Science Foundation Graduate Research Fellowship Program (NSF GRFP) under Grant No. 2141064. J.H. acknowledges support from MIT Thomas and Sarah Kailath Fellowship.

## Author contributions

K.L., L.G.G., E.C., and S.Y. conceived the idea of the project. K.L. built the optical setup and performed the imaging experiments with critical inputs from H.C. K.L. designed the network training scheme and performed implementations with critical inputs from J.H.. E.L.K. and E.C. designed and performed the biological experiments for imaging and characterization. K.L. and E.C. performed antibody staining and confocal imaging. G.D.N. performed CLEM imaging. K.L. and E.L.K. analyzed the data and prepared the figures. K.L., E.L.K., E.C., and S.Y. wrote the manuscript with input from all authors. S.Y., E.C., and L.G.G. obtained the funding and supervised the research.

